# Minimum inoculum of resistance assay for evaluating for anti-toxoplasmosis compounds that target phenyl alanine tRNA synthetase

**DOI:** 10.64898/2026.03.02.709153

**Authors:** Taher Uddin, Payal Mittal, Han Xie, Bruno Melillo, Amit Sharma, Arnab K. Chatterjee, L. David Sibley

## Abstract

*Toxoplasma gondii* is a globally important intracellular parasite, and treatment regimens are limited by the failure of drugs to target latent tissue cysts. Developing new candidates for treatment also needs to address the potential for resistance to arise. Here, we developed a Minimum Inoculum for Resistance (MIR) assay as a quantitative metric for evaluating inhibitors of *T. gondii*. The MIR assay, adapted from assays used in malaria drug discovery, measures the frequency for pre-existing resistance alleles by exposing different sized parasite populations to drug pressure. We profiled a series of bicyclic pyrrolidone analogs that inhibit phenylalanine tRNA synthetase (PheRS). We demonstrate that these inhibitors require higher inocula to lead to parasite resistance (up to > 10^8^ parasites) in comparison with an inhibitor of DNA synthesis, and that MIR values vary across inhibitors with closely related chemical structures. Clonal analysis of resistant parasites emerging from MIR assays revealed both new and previously identified resistance conferring mutations in *Tg*PheRS, and structural modeling revealed their potential impact the enzyme active site. The MIR assay provides a functional benchmark to compare new and existing inhibitors, allowing for rational prioritization of lead compounds with a high genetic barrier to resistance.

## Introduction

*Toxoplasma gondii* is an obligate intracellular protozoan parasite responsible for toxoplasmosis, a globally prevalent infection with significant human and veterinary health impacts ^1-5^. *T. gondii* infection in humans usually results mild symptoms, consisting of fever, aches, and swollen glands, that resolve in weeks in healthy individuals. However, in immunocompromised people like those with HIV/AIDS, infection can cause severe issues affecting the brain, eyes, and lungs ^6-8^. Additionally, in the case of pregnancy, congenital infection can affect development of the fetus and result in defects that range from serious CNS deformations to mild cognitive impairment and ocular complications later in life ^9^. Persistent infection depends on transition from the rapidly replicating tachyzoite to the slow-growing, semi-dormant bradyzoite within tissue cysts that reside in muscle and brain tissues that reside in muscle and brain tissues and can reactivate, leading to serious illness ^10^. The chronic stages predispose individuals to the potential to reactivation while the actively replicating tachyzoite form is responsible for clinical disease.

Standard therapies such as pyrimethamine and sulfadiazine are effective at stopping *T. gondii* tachyzoite proliferation, but they fail to eliminate the chronic bradyzoite in tissue cysts because bradyzoites are metabolically semi dormant and well-protected within the cyst wall ^11-13^. Additionally, long-term, high-dose therapies can result in severe side effects ^14^. Novel strategies, including combination therapies, cyst wall-targeting drugs, and immunotherapies, are under investigation but require extensive testing for efficacy and safety before they could advance to the clinic ^15^. Antimicrobial resistance is a universal challenge for anti-infective drugs and often reflect heritable, genetically encoded changes in microbes that reduce a drug’s efficacy ^16^. A high barrier to resistance is essential for moving leads into the clinic, thus resistance risk should be assessed with standardized quantitative assays. The minimum inoculum for resistance (MIR) assay, originally established in malaria research, provides a quantitative framework for resistance at the preclinical stage ^17-19^. By determining the smallest population of parasites from which pre-existing resistance can emerge under selective drug pressure, the MIR assay provides a filter for prioritizing clinical candidates. Historical examples in antimalarial drug development have shown that drugs with low MIR values frequently fail in clinical trials due to rapid resistance emergence, while compounds with high MIR are more likely to show durable efficacy and become standard therapies ^17, 20-23^. Thus, MIR data can also influence lead advancement by allowing for deprioritization of molecules with unacceptable resistance risk.

In this study, we established an MIR protocol for *T. gondii* that defines inoculum thresholds for resistance. The MIR assay in *Toxoplasma* research will facilitate prioritization of compounds with high genetic barriers to resistance, and thus speed discovery of durable therapies for toxoplasmosis.

## Results and Discussion

To establish a MIR assay for *T. gondii*, we systematically exposed cultures infected with tachyzoites to high-level, single-step drug pressure. For these assays, we used human foreskin fibroblast (HFF) cultures seeded with different parasite inocula (10^6^–10^8^ *Tg*Me49-Fluc tachyzoites) and treated them with 3X EC_90_ concentrations of candidate compounds in triplicate populations (defined as F1, F2, F3) for 30 days (Figure 1A, 2A-E). We emplyed a low-passage, clonal *Tg*Me49-Fluc parasite line B6 that was isolated by limited dilution using standard procedures. We compared the resistance profile of previously characterized potent and selective inhibitors of *T. gondii* phenylalanyl-tRNA synthetase (PheRS) including one 4,8-bicyclic azetidine compound (referred to as BRD7929) and six 5,7-bicyclic pyrrolidine analogues ^24-26^ (Figure 1C, Table S1). As a positive control, we used 5-fluoro-2’-deoxyuridine (FUDR), which is a substrate analog incorporated into fluorodeoxy-UMP by the salvage enzyme uracil phosphoribosyl transferase (UPRT), thus blocking the activity of thymidylate synthetase (Figure 1C). The primary mechanism for FUDR resistance in *T. gondii* is the loss of UPRT activity; when it is inactivated, the parasite cannot convert FUDR into toxic metabolites thus allowing survival ^27, 28^. As anticipated, robust resistance to FUDR emerged rapidly within two weeks at lowest inoculum tested (<10^6^) (Figure 1B, 2A). In contrast, resistance to test compounds were observed only under higher inoculum pressures. For example, selection with compound **1** (previously referred to as BRD2108) and compound **10** resulted in resistant lines at an inoculum of 10^7^ in all three populations (F1, F2, F3), while compound **21** required an inoculum of 10^8^ to generate resistant population in all three populations (F1, F2, F3). In contrast, compound **15** selection resulted in resistance in one (F3) out of three flasks/population at an inoculum of 10^8^ (Figure 1B, 2B-E). No resistance to compounds **12, 19**, or BRD7929 was observed up to an inoculum of 10^8^, which was the maximum level tested here (Figure 1B). Despite the close structural similarity of these compounds that share the same PheRS target ^24-26^, they varied substantially in the different inoculum sizes required to achieve resistance.

**Figure 1:**
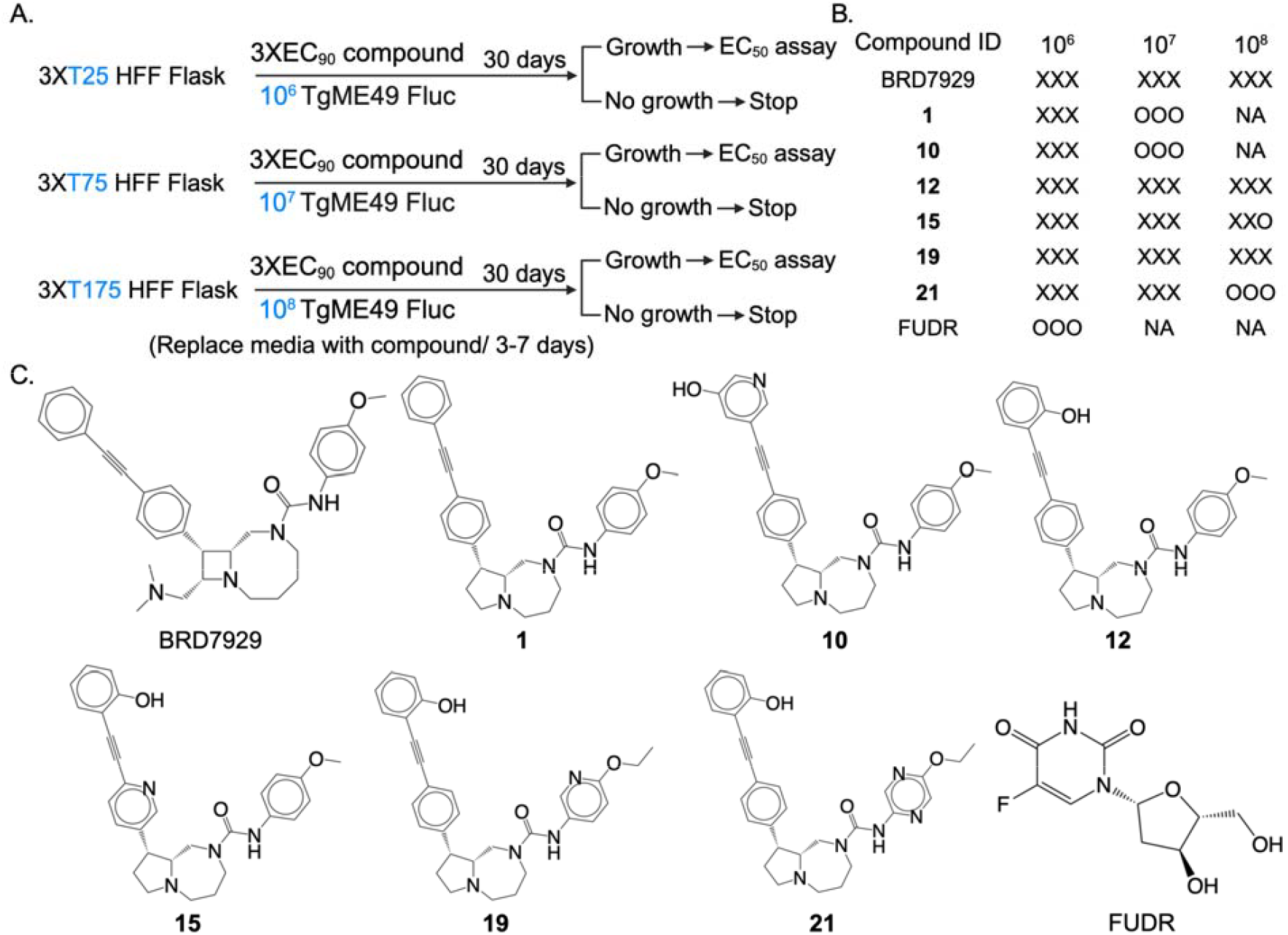
Schematic overview of the Minimum Inoculum for Resistance (MIR) assay and inhibitors used in this study. A. MIR assay workflow: confluent HFF cultures were seeded with different parasite inocula (10^6^ to 10^8^) and exposed to 3XEC_90_ concentrations of each compound. Flasks were monitored for 30 days, with compound-containing media refreshed every 3–7 days. Upon parasite growth, parasites were sub-cultured, and EC_50_ values were assessed via luciferase assay. B. Schematic summary of outcomes for compounds tested in this study: X = No parasite growth in 30 days. Y = parasite growth in 30 days. Each X or Y represents one flask. NA-Not applicable. C. Chemical structures of the *Tg*cPheRS inhibitors and thymidylate synthase inhibitor (FUDR) evaluated in the MIR assay.

In cases where resistant parasites emerged in the culture, we evaluated the resistance profile using *in vitro* luciferase-based growth to estimate the EC_50_. Parasites resistant to FUDR were observed at an inoculum of 10^6^ in a similar timeline in all three populations; however, separate populations (F1, F2, F3) showed different fold resistance (FR) calculated by the difference in EC_50_ between resistant lines and the parental Me49Fluc line (Figure 2A). Compound **1** resistant populations also showed different levels of fold resistance in at an inoculum of 10^7^ (Figure 2B). In contrast, three compound **10** resistant populations showed lower-level fold resistance in at an inoculum of 10^7^ (Figure 2C). The sole compound **15** resistant population (F3) that developed at an inoculum of 10^8^ had a FR of 3 (Figure 2D). Like compound **1**, compound **21** resistant populations in that developed at an inoculum of 10^8^ also showed diverse fold resistance (Figure 2E).

**Figure 2:**
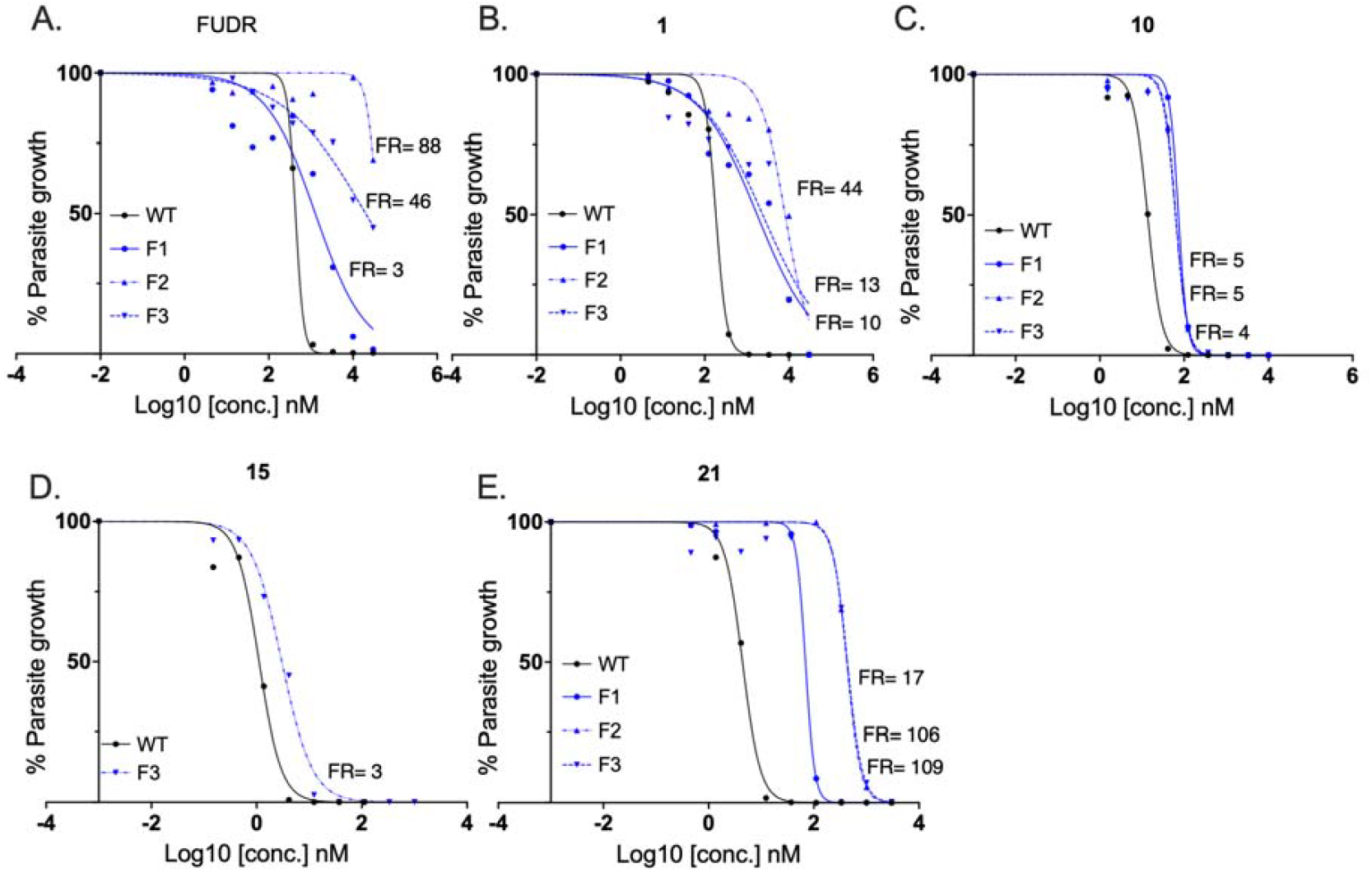
EC_50_ inhibition curves. A. WT vs three FUDR resistant parasite populations F1, F2, F3. B. WT vs three Compound **1** resistant parasite populations F1, F2, F3. C. WT vs three compound **10** resistant parasite populations F1, F2, F3. D. WT vs one compound **15** resistant parasite population F3. E. WT vs three compound **21** resistant parasite populations F1, F2, F3. All EC_50_ values are presented as the mean of two technical replicates. Source data are provided in Table S3. FR= fold resistance; WT= parental TgMe49Fluc parasite.

To evaluate the clonal populations further, single cell clones were generated by limiting dilution from all parasite populations resistant to the compound **1, 10, 15** and **21**. Two to three clones (named as C1, C2, C3) per population were cultured to evaluate fold resistance in the individual lines (Table1). With the exceptions of compound **10** F2C2 and compound **15** F3C3 clonal lines, the fold resistance in clonal parasite lines mirrored that seen in the population (Table1). We also tested the sensitivity of clonal parasite lines to the compounds FUDR, atovaquone and pyrimethamine to evaluate the potential for cross resistance. None of the resistant parasite lines showed increased resistance to these compounds (Table 1).

**Table 1:** Fold resistance and mutation profile of population and clonal parasite lines.

| Compound ID<br>Inocula size | Population<br>(EC<sub>50</sub> FR- self) | Clone | EC<sub>50</sub> FR<br>(self) | EC<sub>50</sub> FR<br>(FUDR) | EC<sub>50</sub> FR<br>(ATQ) | EC<sub>50</sub> FR<br>(PyR) | TgPheRS<br>mutation |
| --- | --- | --- | --- | --- | --- | --- | --- |
| **1**<br>10<sup>7</sup> | F1 (10) | C1 | 14 | 2 | 1 | 1 | L425V |
|  |  | C2 | 15 | 2 | 1 | 2 | L425V |
|  |  | C3 | 11 | 2 | 1 | 1 | L425V, L521P |
|  | F2 (44) | C1 | 63 | 1 | 1 | 1 | L497V |
|  |  | C2 | 39 | 1 | 1 | 1 | L497V |
|  | F3 (13) | C1 | 43 | 1 | 1 | 1 | L497V |
|  |  | C2 | 42 | 1 | 1 | 1 | L497V |
| **10**<br>10<sup>7</sup> | F1 (5) | C1 | 5 | 1 | 2 | 1 | M260I |
|  |  | C2 | 4 | 2 | 2 | 2 | M260I |
|  | F2 (5) | C1 | 6 | 2 | 2 | 1 | M260I |
|  |  | C2 | 86 | 1 | 1 | 1 | M260I, W495C |
|  | F3 (4) | C1 | 4 | 1 | 1 | 1 | M260I |
|  |  | C2 | 3 | 1 | 1 | 1 | M260I |
| **15**<br>10<sup>8</sup> | F3 (17) | C1 | 4 | 2 | 1 | 1 | L497V |
|  |  | C2 | 3 | 1 | 1 | 1 | L497V |
|  |  | C3 | 100 | 1 | 2 | 1 | L497V, W495L |
| **21**<br>10<sup>8</sup> | F1 (17) | C1 | 5 | 2 | 1 | 1 | V262A |
|  |  | C2 | 4 | 2 | 1 | 1 | V262A |
|  | F2 (106) | C1 | 99 | 2 | 2 | 1 | L497V |
|  |  | C2 | 85 | 2 | 2 | 1 | L497V |
|  | F3 (109) | C1 | 43 | 1 | 1 | 1 | L497V |
|  |  | C2 | 63 | 1 | 1 | 1 | L497V |

F=(Flask/population), C=clone, FR = EC<sub>50</sub> fold resistance between resistant parasite line and parental TgMe49 parasite line. FUDR = 5-fluoro-2'-deoxyuridine, ATQ = Atovaquone, PyR = Pyrimethamine. MIR= Minimum Inoculum for Resistance. TgPheRS = *Toxoplasma gondii* phenylalanyl-tRNA synthetase. self = tested against the resistance-inducing compound.

To determine the molecular basis of resistance, genomic DNA from the parasite clonal lines were used to amplify the presumed target of the compounds, the cytosolic α subunit of *Tg*PheRS (TGME49_234505, toxodb.org) followed by next generation sequencing to define potential mutations (Table 1). Two of the clones (C1, C2) from compound **1** selected F1 population carried a L425V mutation, whereas the 3^rd^ clone (C3) carried an additional mutation L521P in addition to L425V (Table 1). Two clones (C1, C2) from other two compound **1** selected F2, F3 population carried a L497V mutation (Table 1). Two clonal lines (C1, C2) from each compound **10**-resistant population (F1, F2, F3) carried a M260I mutation. Only one of the clones (compound **10** F2C2) carried an additional W495C mutation (Table 1). For compound **15**, the resistant clones C1 and C2 carried L497V mutations, whereas C3 carried an additional W495L mutation in additional to L497V (Table 1). For compound **21** resistant parasites, two clones from F1C1 and F1C2 showed a mutation V262A, and four clones F2C1, F2C3, F3C1, F3C2 exhibited a L497V mutation (Table 1). Next, we tested all clonal parasite lines for cross resistance to the same panel of six compounds (Figure 3). Clonal lines that showed high resistance to their selecting compound also showed high resistance to other compounds of the panel, whereas lines with low resistance to their parental compound showed correspondingly low cross resistance (Figure 3).

**Figure 3:**
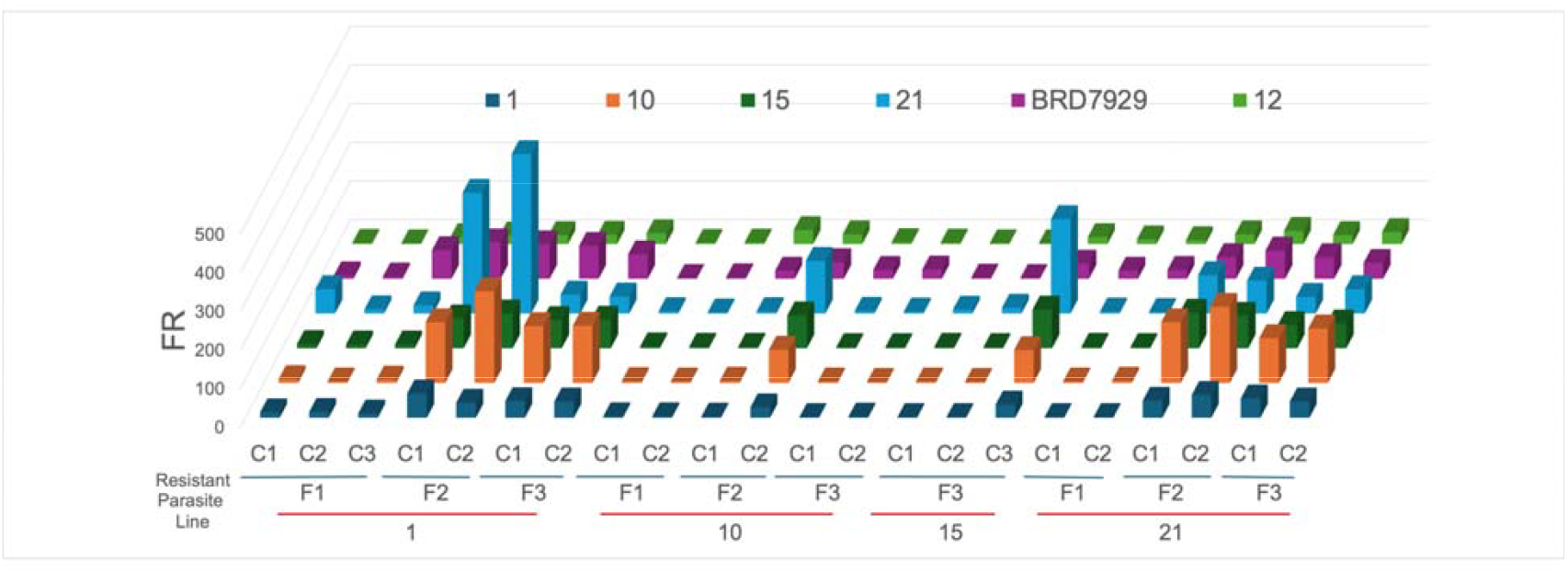
Cross-resistance profiles of clonal resistant parasite lines to *Tg*cPheRS inhibitors compounds. The X-axis represents individual clonal resistant parasite lines generated under selective pressure with each of four compounds. The Y-axis denotes the small molecule inhibitors depicted in different colors. The Z-axis indicates the fold resistance (EC_50_ of clonal resistant parasite/parental parasite). F=(Flask/population), C=clone, FR = EC_50_ fold resistance between resistant parasite line and the parental TgMe49Fluc parasite line.

Prior work reported selection of BRD7929-resistant *T. gondii* Me49 parasites by cultivation in gradually increasing concentrations over 5 months, resulting in 10-to 100-fold EC_50_ elevations ^26^. These studies identified mutations in the *Tg*PheRS α-subunit mutations (L497V/I, V262A, M484I) as principal resistance determinants ^26^. Introduction of the L497I mutation alone into a wild-type background conferred strong resistance (40–60-fold EC_50_ shift), while V262A and M484I mutations contributed lower levels of resistance (2–4-fold) validating *Tg*PheRS as the primary resistance locus ^26^. Our study expanded on this prior work by testing the existence of propensity of resistance to arise within a defined starting inoculum selected at a single high concentration. Like prior studies, our findings indicate that most clones that carry a mutation at L497V show high resistance, while mutations at V262A lead to low resistance. Somewhat surprisingly, two clones carrying L497V (compound **15** F3, C1, C2) showed only low-level resistance, despite the fact that previous studies have shown that introduction of this single change is sufficient to confer high level resistance ^26^. We retested the sequence of the *Tg*PheRS α-subunit and the resistance profile of these clones and confirmed the original finding. In addition, we also checked the *Tg*PheRS β-subunit (TGME49_306960), however did not identify any mutation. Although we have not defined the basis for the low-level resistance of clones harboring L497V here, it suggests an unknown extragenic change that muted the normal resistance profile of this mutation. Additionally, we identified four new mutations (M260I, L425V, W495L, L521P) (Table 1). Clones with the L425V mutation showed moderate resistance, while those harboring M260I showed low resistance, with the exception for compound **10** clone F1, C2, which also harbored a M495C mutation leading to high resistance (Table 1).

To gain more insight into the molecular basis of resistance, we generated a structural model of *Tg*PheRS using AlphaFold3, as no crystallographic structure for the inhibitor-bound enzyme is currently available (Figure 4A). The model was of high quality, with a pLDDT score exceeding 90 in the catalytic domain and consistent inter-domain confidence (iPTM = 0.68), allowing reliable interpretation of mutation impacts. After energy minimization and hydrogen bond optimization, the aminoacylation active site and substrate-binding pockets were annotated using homology with known FRS structures. Docking studies were then performed with representative bicyclic pyrrolidine analogs using the Glide XP protocol, with the best-ranked poses retained for analysis. We mapped resistance-associated mutations (V262A, M260I, L425V, M484I, L497I/V, W495L, L521P) onto the *Tg*PheRS–ligand complexes using UCSF Chimera, differentiating those newly identified in this study from previously reported sites (Figure 4B). This approach revealed that novel low resistance mutations, such as L425V, M260I, and L521P localize outside of the ligand-binding region, whereas high resistant mutation W495L located near the ligand-binding region, suggesting potential mechanisms for altered inhibitor affinity or substrate competition. The model also supports the functional importance of residues such as V262 and L497 which correspond to resistance hot spots in related apicomplexan PheRS enzymes ^26^. These findings provide a structural and mechanistic framework for understanding the diverse mutation profiles observed in MIR-selected resistant clones.

**Figure 4.**
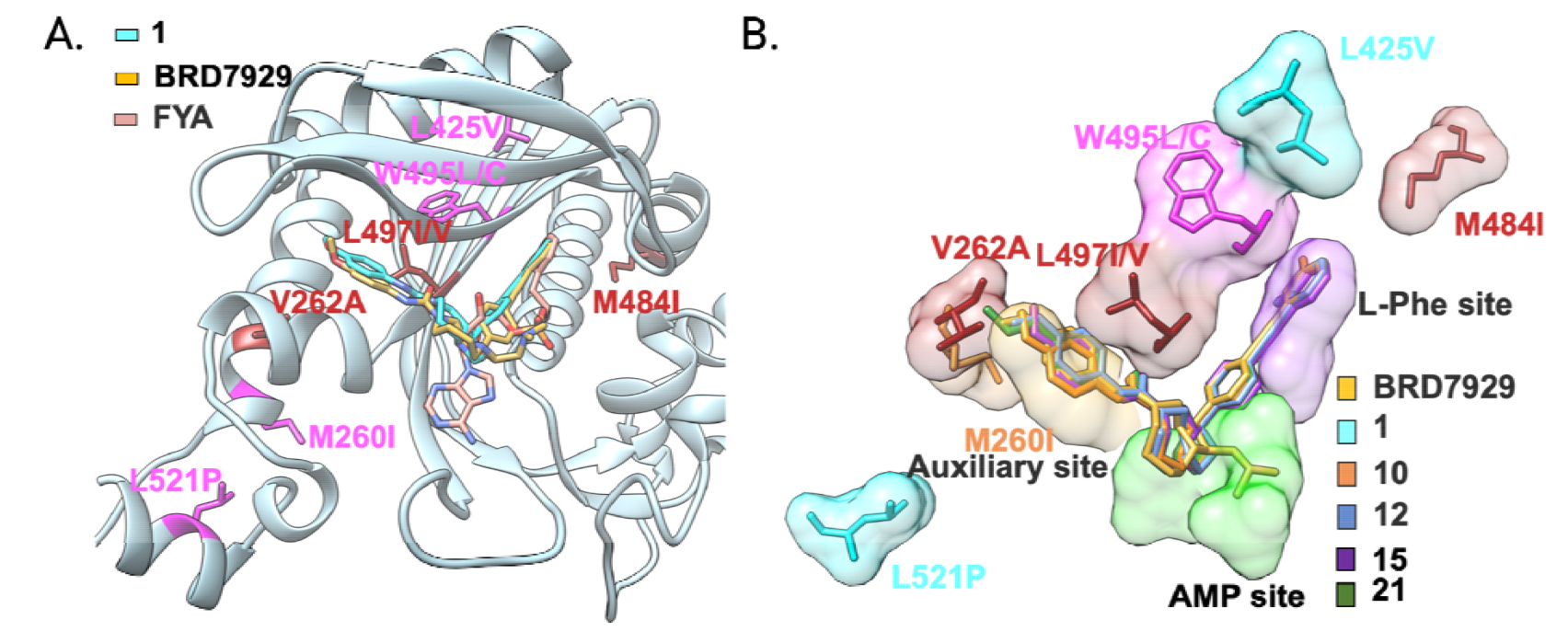
Mutation mapping in *Tg*PheRS. A. Homology model of the alpha-subunit of *Tg*cPheRS (light blue) docked with compound **1** shown in cyan, phenylalanyl-adenylate analogue in salmon (FYA), and BRD7929 in gold. The drug-resistant *Tgc*PheRS mutations found previously with BRD7929 are highlighted in red ^26^. The new mutations found in this study are highlighted in pink. B. Superposition of docked bicyclic pyrrolidines within the alpha-subunit of *Tg*cPheRS enzyme. The *E. coli* PheRS bound with AMP and L-Phe (PDB ID: 3PCO) was superimposed to delineate active site pockets. The AMP pocket is shown in green, the L-phenylalanine pocket in light purple and the auxiliary site in gold. Mutations associated with resistance are indicated along with their corresponding compounds. Previously described mutations (dark red) include V262A, M484I, and L497I/V with BRD7929. New mutations include M260I (orange), L425V, L521P (cyan) and W495L (magenta).

## Conclusion

The MIR assay described here evaluates the potential for resistance to either be pre-existing or rapidly emergent under strong selection. Under these conditions, we observed a similar pattern of mutations that give rise to high-level resistance when compared to previous studies that used stepwise selection with increasing concentrations of compound. In addition, we identified other potential sites of low-level resistance. Mapping these onto the structure of *Tg*PheRS reveals a general pattern of high-level resistance being due to changes in the active site, while changes at more distant sites led to lower-level resistance. Surprisingly, we observed widely different propensity for the rapid emergence of resistance using structurally related compounds that target the *Tg*PheRS enzyme. The different frequencies of resistance where not correlated with the relative potency, selectivity, hydrophobicity, or other physical-chemical properties of the chosen compounds. Although we cannot rule out that some compounds are protected from resistance by unknown secondary targets, these studies are important for illustrating that the MIR for a given compound class can vary with subtle structural variants of the scaffold.

In summary, this study establishes the MIR assay as a quantitative resistance-risk framework for *T. gondii* and demonstrates its utility for distinguishing compounds with varying propensities to drive resistance emergence. Adoption of MIR-based resistance profiling will strengthen preclinical prioritization strategies and accelerate the development of next-generation anti-toxoplasmosis therapies with improved long-term efficacy.

**Table S1:** Potency, *Tg*L497I resistance profile and host cell line selectivity of 270 MIR Study candidate compounds ^24^.

| Compd ID | *Tg*Me49<sup>a</sup><br>EC<sub>50</sub> | *Tg*Me49<br>EC<sub>90</sub> | *Tg*L497I EC<sub>50</sub>/<br>Parental<sup>b</sup> EC<sub>50</sub> (FR)<sup>c</sup> | HepG2 CC<sub>50</sub> | SI<sup>d</sup> | THP-1 CC<sub>50</sub> | SI<sup>e</sup> |
| --- | --- | --- | --- | --- | --- | --- | --- |
| BRD7929 | 0.048 ± 0.026 | 0.132 ± 0.082 | 78 | 6.04 ± 1.11 | 147 | 1.01 ± 0.03 | 24 |
| **1** | 0.291 ± 0.034 | 0.439 ± 0.043 | 10 | 18.45 ± 0.67 | 51 | 14.5 ± 0.133 | 48 |
| **10** | 0.011 ± 0.003 | 0.034 ± 0.001 | 100 | 38.70 ± 11.77 | 2056 | 6.42 ± 2.77 | 385 |
| **12** | 0.001 ± 0.000 | 0.002 ± 0.001 | 47 | 1.12 ± 0.27 | 1447 | 0.18 ± 0.12 | 374 |
| **15** | 0.003 ± 0.008 | 0.004 ± 0.001 | 36 | 4.66 ± 1.43 | 1462 | 0.51 ± 0.30 | 196 |
| **19** | 0.003 ± 0.001 | 0.006 ± 0.002 | 50 | 1.18 ± 0.22 | 369 | 0.21 ± 0.08 | 73 |
| **21** | 0.007 ± 0.001 | 0.016 ± 0.001 | 28 | 4.16 ± 0.29 | 797 | 2.72 ± 2.55 | 402 |
| FUDR | 0.499 ± 0.112 | 0.643 ± 0.286 | nd | nd | na | nd | na |

Values in μM, average of three or more biological replicates. <sup>a</sup> *Tg*Me49 expressing firefly luciferase, <sup>b</sup> Parental strain used is *Tg*RHΔ<sub>hxgprt</sub>Δ<sub>ku80</sub> (*Tg*RHΔΔ), <sup>c</sup> EC<sub>50</sub> fold resistance between BRD7929 resistant parasite line *Tg*L497I and parental parasite line *Tg*RHΔΔ. <sup>d</sup> Selectivity Index (HepG2 CC<sub>50</sub>/*Tg*Me49EC<sub>50</sub>); <sup>e</sup> Selectivity Index (THP-1 CC<sub>50</sub>/*Tg*Me49EC<sub>50</sub>) nd = not determined. na=not applicable

**Table S2:**
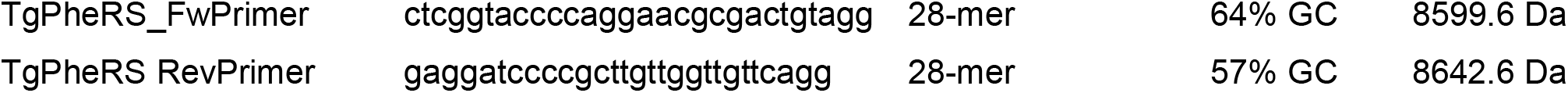
Primer sequences used to amplify the TgPheRS.

**Table S3:**
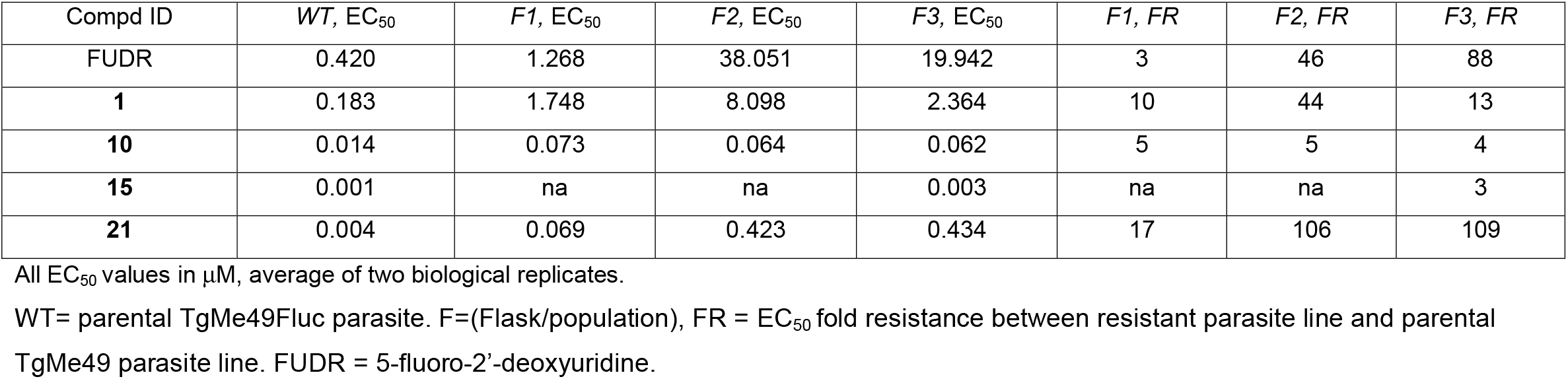
Potency and fold resistance profile of parasite populations.

| Compd ID | WT, EC<sub>50</sub> | F1, EC<sub>50</sub> | F2, EC<sub>50</sub> | F3, EC<sub>50</sub> | F1, FR | F2, FR | F3, FR |
| --- | --- | --- | --- | --- | --- | --- | --- |
| FUDR | 0.420 | 1.268 | 38.051 | 19.942 | 3 | 46 | 88 |
| **1** | 0.183 | 1.748 | 8.098 | 2.364 | 10 | 44 | 13 |
| **10** | 0.014 | 0.073 | 0.064 | 0.062 | 5 | 5 | 4 |
| **15** | 0.001 | na | na | 0.003 | na | na | 3 |
| **21** | 0.004 | 0.069 | 0.423 | 0.434 | 17 | 106 | 109 |

All EC<sub>50</sub> values in $\mu$ M, average of two biological replicates.
WT= parental TgMe49Fluc parasite. F=(Flask/population), FR = EC<sub>50</sub> fold resistance between resistant parasite line and parental
TgMe49 parasite line. FUDR = 5-fluoro-2'-deoxyuridine.

## Experimental Methods

### Compound source and *in vitro* luciferase-based potency evaluation

Compounds were obtained from Calibr (Scripps Research, La Jolla, CA 92037, USA) either as lyophilized powder or 10 mM stocks in 100% DMSO and stored at −80°C. Human foreskin fibroblast (HFF) cell line and parasites were cultured in a medium consisting of 10% DMEM (Dulbecco’s Modified Eagle Medium, D10), supplemented with 10 mM glutamine, 10 μg/mL gentamycin, and 10% fetal bovine serum. The cultures were incubated at 37°C with 5% CO_2_ and routinely confirmed negative for mycoplasma contamination using the e-Myco Plus kit from Intron Biotechnology. Luciferase-based growth assays were conducted using the previously described TgMe49-Fluc strain ^29^. To reduce variability, only the inner 60 wells of white 96-well plates (Costar) were used to generate a 3-fold series of compounds. Each well received 10×10^3^ freshly harvested TgMe49-Fluc or resistant lines tachyzoites cultured in confluent HFF in 100 µL volumes, achieving a final compound concentration of 200 µL/well with 0.1% DMSO in 10% DMEM. Plates were incubated for 72 h at 37°C with 5% CO_2_ before analysis, following the Promega Luciferase Assay System protocol. After incubation, culture media were replaced with 50 µL of 1x Cell Culture Lysis Reagent (Promega). After 10 minutes at room temperature, 100 µL of Luciferase Assay reagent (Promega) was added using an injection system, and luciferase activity was measured using a BioTek Cytation 3 multi-mode imager with Gen5 software.

### Generation of compound resistant parasite lines and MIR assay

To evaluate the MIR, 8 compounds, including 7 previously reported^24-26^ and 5-Fluorodeoxyuridine (FUDR), were used in this study. Potency, selectivity, and fold resistance to previously reported BRD7929-specific point-mutated TgRH-PheRS^[L497I]^-FLuc parasite line ^24-26^are summarized in Table S1, modified from previously published data ^24-26^. Triplicate HFF confluent T25 flasks were prepared in D10 media for each compound. Freshly harvested 10^6^ TgMe49-Fluc tachyzoites (MIR 10^6^) were added to each flask, with the D10 media supplemented to a 3XEC_90_ concentration of the compound (Figure 1A). The flasks were monitored for 30 days, with media replaced by fresh compound-supplemented D10 media every 3-7 days. If parasite growth was observed under microscope, parasites were subculture, and TgEC_50_ values were assessed using the luciferase assay protocol described above. In parallel, triplicate HFF confluent T75 flasks with 10^7^ TgMe49-Fluc tachyzoites (MIR 10^7^), and triplicate HFF confluent T175 flasks with 10^8^ TgMe49-Fluc tachyzoites (MIR 10^8^) were set up and followed up similarly (Fig 1A). This parallel approach systematically evaluated the development of compound-resistant parasite lines with requirement of minimum inoculum of parasite. Resistance detected at any stage prompted cloning resistant parasites via minimum dilution in 96-well plates. TgEC_50_ and fold differences between at least two clonal resistant lines and the parental TgMe49-Fluc line (WT) were evaluated as described above, and gDNA from each clonal line were extracted for PCR amplification of *Tg*PheRS α-subunit (Toxodb.org, TGME49_234505) and β-subunit (TGME49_306960), and sequence for genetic mutations.

### gDNA extraction, PCR, Sequencing, Mutation mapping

Infected HFF cells were scraped and syringe-lysed using 25 g needles to release the parasites. The parasites were harvested by passage through a 3-μm filter and centrifugation at 400 × g for 10 min. gDNA was isolated using DNeasy Blood & Tissue Kit (QIAGEN). TgPheRS sequence was amplified using PrimeSTAR GXL Premix (Takara bio), PCR primer sequences are provided in Table S2. After gel conformation and cleaning the PCR product using PCR cleanup and gel extraction kits (Takara bio), plasmidsaurus sequence amplicons service that utilizes the ONT Rapid Barcoding Kit V14 and is sequenced on a PromethION flow cell to get full length coverage of PCR products, with no primers. Sequence data was analyzed using SnapGen 8.2.0.

### Structure-models of the resistance mutations

As the crystallographic structure of *T. gondii* phenylalanyl-tRNA synthetase (*Tg*cPheRS) in complex with inhibitors is not yet deposited in the Protein Data Bank (PDB), the *Tg*cPheRS model was generated using AlphaFold3^24 30^ based on the full-length amino acid sequence of the alpha-subunit ( Toxodb.org, TGME49_234505). The predicted inter-domain predicted TM (iPTM) is 0.68, which indicates high overall model confidence and a very high pLDDT score (>90) for most of the residues in the catalytic domain. The model was energy-minimized in Schrödinger’s Protein Preparation Wizard (Schrödinger Release Maestro version 14.2.121, MMshare Version 6.8.121, Release 2024-4, Platform Darwin-x86_64) with default parameters, ensuring correct protonation states at pH 7.4 and optimization of the hydrogen-bonding network. The binding site was defined based on the aminoacylation active site and phenylalanine/ATP binding pockets inferred from homologous FRS structures. Compounds were prepared using LigPrep with the OPLS4 force field and all the possible states were determined at target ph 7.5+/-1.00 and specific chirality were retained. Molecular docking was performed using Glide XP mode. The best-ranked poses for each compound were retained for visualization. Resistance-associated mutations (L425V, W495L, V262A, M260I, L497I/V, M484I) were mapped onto the *Tg*cPheRS–ligand complexes in UCSF Chimera v1.16, with residues color-coded to distinguish newly identified mutations from those previously reported.

### Statistical Analysis

Statistical analyses were performed using Prism 10 (GraphPad Software, Inc.). Dose-response inhibition curves for parasite and host cell toxicity screens (EC_50_ and CC_50_ values) were generated using the “Log(inhibitor) vs. normalized response—Variable slope” function.

## Author contributions

Conceptualization: T.U., L.D.S; Data curation: T.U.; Formal analyses: T.U., P.M.; Investigation: T.U., P.M.; Supervision: L.D.S.; Writing original draft: T.U., P.M.; Writing review and editing: all authors.

## Acknowledgements

We thank Jennifer Bark for assistance with cell culture. We thank Laura Knoll for providing the TgME49Fluc strain. Supported in part by a grant from the National Institutes of Health (AI143857).

